# Conservation of single amino-acid polymorphisms in *Plasmodium falciparum* erythrocyte membrane protein 1 and association with severe pathophysiology

**DOI:** 10.1101/128983

**Authors:** Daniel Zinder, Mary M. Rorick, Kathryn E. Tiedje, Shazia Ruybal-Pesántez, Karen P. Day, Mercedes Pascual

## Abstract

*Plasmodium falciparum* erythrocyte membrane protein 1 *(PfEMP1)* is a parasite protein encoded by a multigene family known as *var.* Expressed on the surface of infected red blood cells, *PfEMP1* plays a central role in parasite virulence. The *DBLα* domain of *PfEMP1* contains short sequence motifs termed homology blocks. Variation within homology blocks, at the level of single amino-acid modifications, has not been considered before in association with severe disease. Here we identify a total of 2701 amino-acid polymorphisms within *DBLα* homology blocks, the majority of which are shared between two geographically distant study populations in existing transcription data from Kenya and in a new genomic dataset sampled in Ghana. Parasitemia levels and the transcription levels of specific polymorphisms are as predictive of severe disease (AUC=0.83) and of the degree of rosetting (forecast skill SS=0.45) as the transcription of classic *var* groups. 11 newly categorized polymorphisms were strongly correlated with *grpA var* gene expression (SS=0.93) and a different set of 16 polymorphisms was associated with the *H3* subset (SS=0.20). These associations provide the basis for a novel method of relating pathophysiology to parasite gene expression levels—one that, being site-specific, has more molecular detail than previous models based on *var* groups or homology blocks. This newly described variation influences disease outcome, and can help develop anti-malarial intervention strategies such as vaccines that target severe disease. Further replication of this analysis in geographically disparate populations and for larger sample sizes can help improve the identification of the molecular causes of severe disease.

## INTRODUCTION

The manifestation of *Plasmodium falciparum* infection is wildly variable. In highly endemic areas the vast majority of infections are asymptomatic and most cases of clinical malaria are mild. Only a small fraction of individuals with clinical malaria, predominantly infants, go on to develop life-threatening severe disease (1, 2). Severe cases represent less than one percent of infected individuals, but they are nevertheless the dominant cause of morbidity and mortality from this disease (3–5). Despite decades of intensive control efforts, there are approximately 212 million cases of malaria annually, and hundreds of thousands of deaths due to *P. falciparum* infection, the majority of which are children in Africa (6). The fact that most people in endemic regions rapidly develop immunity to complicated malaria within the first few years of life offers hope for the possibility of developing targeted vaccines to prevent severe disease (7).

In the bloodstream, *P. falciparum* infects red blood cells and then uses a diversity of human receptors to promote the adherence and sequestration of infected red blood cells (iRBCs) within various human tissues. Parasite density, the fraction of iRBCs, is associated with increased virulence, and *P. falciparum’s* unique ability to readily invade mature RBCs contributes to its increased pathogenicity (8). While a high parasite burden is more common during severe disease, improved control of parasite density alone does not explain resistance to severe disease (9). iRBCs avoid clearance by the spleen by tethering to the linings of small blood vessels in various host tissues, and by binding uninfected cells—a phenomenon called rosetting. Massive iRBC sequestration in host microvasculature strongly contributes to virulence (10). Some severe complications of malaria are associated with parasite sequestration in particular tissues: e.g. in the microvasculature of the brain in the case of cerebral malaria, and the placenta in the case of pregnancy-associated malaria. Other severe pathophysiologies include extreme weakness, convulsions, renal failure, circulatory shock, low blood glucose levels, acidosis, respiratory distress, impaired consciousness and severe malarial anemia (11).

The adherence and sequestration of the parasite in particular tissue types, as well as the rosetting phenotype, are strongly associated with the expression of specific members of the *var* gene family (12, 13). The *var* genes encode *Plasmodium falciparum* erythrocyte membrane protein 1 (P*f*EMP1), and specific P*f*EMP1 types have been found to be preferentially expressed in particular host tissues including the brain and cardiac tissue of severely diseased infants and in infected placental tissue (12, 14–16). Located in multiple *subtelomeric* and central chromosomal regions, approximately 60 antigenically distinct P*f*EMP1 variants are encoded per parasite genome. P*f*EMP1 proteins are large (200-350 kDa) and come in a diversity of architectural types, which are defined by the presence and multiplicity of specific domains (17). The major architectural subtypes can be classified into three groups—A, B and C, and they are associated with distinct chromosomal locations, transcription direction, and semi-conserved upstream sequence (*ups*) tags (18–20). Because group A/B/C *var* genes are commonly classified by their *ups* tags, and because we do not have *ups* sequences for the *var* diversity considered here, we will use “grpA-like” to refer to a group of *var* gene sequences classified by an alternative network-based method previously established to be highly correlated with *ups*-based group A classification (13, 21).

Because of its important role in binding parasite-infected cells to host microvasculature, the extracellular domains of P*f*EMP1 are under strong natural selection for affinity to host endothelial receptors. Residing in high density on the outside of infected cells, these domains are also under strong natural selection to evade the adaptive immune system. The parasite employs an antigenic variation system whereby individual parasites only express a single *var* gene at a time, switching expression over the course of an infection (8, 22, 23). The regulation of *var* gene expression is not yet fully elucidated, though it appears to centrally involve epigenetic machinery (24–26). As a probable outcome of immune evasion, *var* gene sequences are highly diverse both within individual parasite genomes and at the population level. Within-domain amino acid identity is less than 50%, even within the same architectural type (20, 27). Although sterilizing immunity never develops against *P. falciparum,* the severity of disease is rapidly reduced with repeated infection (7, 28, 29). A component of naturally acquired immunity to disease complications appears to involve an antibody response to PfEMP1 (30–32). As such, identifying conserved epitopes of PfEMP1 which are associated with severe disease is of major interest (23, 33, 34).

Detailed subtyping has enabled the identification of domains and domain cassettes which bind to specific endothelial tissue receptors and to placental tissue (20, 35–41). The head structure of the PfEMP1 protein contains two domains referred to as CIDR and DBLα. Increased transcription of group A *vars* has been linked to rosetting and to a higher likelihood of severe disease (21, 42–45). This is possibly the result of the ability of head structure variants, which are associated with group A *var* types, to preferentially bind endothelial protein C receptor (EPCR) and mediate the formation of rosettes (20, 46). Sequence motifs, such as H3, have also been linked with the rosetting phenotype (47). In addition to the head structure, other domains of PfEMP1 have been implicated in pathology. These include the DBLβ in its binding of ICAM-1 in cerebral malaria and the binding of various domains of the *var2csa* in placental malaria (48, 49). In contrast with group A *vars*, the head domains in group B and C *vars* have been shown to preferentially bind to the host’s CD36 receptor (17, 50). The transcripts of group B *vars* have been identified as more common in symptomatic infection of children with malaria (17).

Host genetics also play a central role in determining the severity of disease. The carrier status of traits such as HbS thalassemia, glucose-6-phosphate dehydrogenase deficiency, complement receptor (CR) 1 deficiency, sickle cell traits and duffy blood groups appears to confer a degree of protection against severe malaria in certain populations (51–56). In addition, in other populations, different ABO blood groups have been shown to reduce rosetting frequency, with group O being the most protective against severe disease (57). Host genetic variation conferring malaria resistance appears to generally be due to recent adaptation (5000-10,000years), vary considerably between human populations and be dependent on parasite genetics (58).

An important advance has been made with the discovery of short conserved sequence motifs or ‘homology blocks’ within the P*f*EMP1 coding sequence, many of which have unique cytoadhesion traits and have been implicated in severe disease. Some have been shown to have preferential expression in patients with severe disease symptoms (e.g., (13). Here we consider variation within homology blocks of the DBLα domain, its conservation across populations, and its relation to severe disease symptoms.

## METHODS

### Study site and sampling of DBLα diversity for the genomic dataset

The “genomic data” used in this study was collected in Bongo District (BD), located in the Upper East Region (UER) of Ghana. Details on the study design, study population and data collection procedures have been described previously (Ruybal-Pesántez et al. 2017). To summarize here: sampling was carried out in two sites (“catchment areas”) of similar human population size, age structure and ethnic composition located ∼10-40 kms apart. Vea/Gowrie was expected to possibly exhibit higher and/or less seasonal transmission than Soe because of its proximity to the Vea dam/irrigation area. The catchment areas were further divided into smaller villages: Vea, Gowrie, Soe Sanabisi and Soe Boko, with participants enrolled from “sections” within these villages (Vea: Gonga and Nayire; Gowrie: Nayire Kura and Tingre; Soe Sanabisi: Tindingo and Akulgoo; and Soe Boko: Tamolinga and Mission Area), meaning that the final dataset contains isolates from one of eight sub-populations in total. All individuals were surveyed in June 2012, which is near the end of the dry season when parasite population sizes and diversity are expected to be at their lowest. Methods related to the microscopy, msp2 PCR and the microsatellite PCR are described in detail in (Ruybal-Pesántez et al. 2017). *Var* DBLα tags were sequenced for 209 parasite positive samples.

### Defining distinct *var* types

Throughout this study we focus on the genetic diversity within the *var* DBLα domain because it is the only domain found in nearly all *var* genes. It is also highly conserved relative to other regions of the *var* gene. For these reasons it is the generally accepted and most popular molecular marker of *var* gene diversity for field-based studies (41, 59–61). For the genetic data, the DBLα sequences were assigned to *var* types using a clustering algorithm in a manner consistent with the commonly used 96% nucleotide identity definition for field studies (62). Each *var* type cluster corresponds roughly to sequences with a >97% amino acid sequence identity. This threshold is consistent with the majority of prior work defining distinct types within DBLα tag sequences because it ensures that each distinct sequence type is very likely to represent a naturally occurring distinct variant, and not merely the result of sequencing errors. Homology blocks and homology block’s polymorphisms were identified within distinct *var* gene types.

### HB polymorphism identification

We translated DNA sequences to AA sequences using the software program EMBOSS Transeq (63, 64). We excluded from the analysis sequences that had an unexpected reading frame, apparent frame shift substitutions or stop codons. Homology blocks (HBs) are conserved units of recombination that are present in *var* genes. Here we only consider those that occur within the DBLα tags of our datasets. HBs were identified using the VARDOM web server (65), with a gathering cut-off of 9.97 to define a match. HBs of a specific type were aligned and all the amino-acid variation in sites containing more than one amino-acid variation were catalogued (Fig. 1).

**Figure 1.**
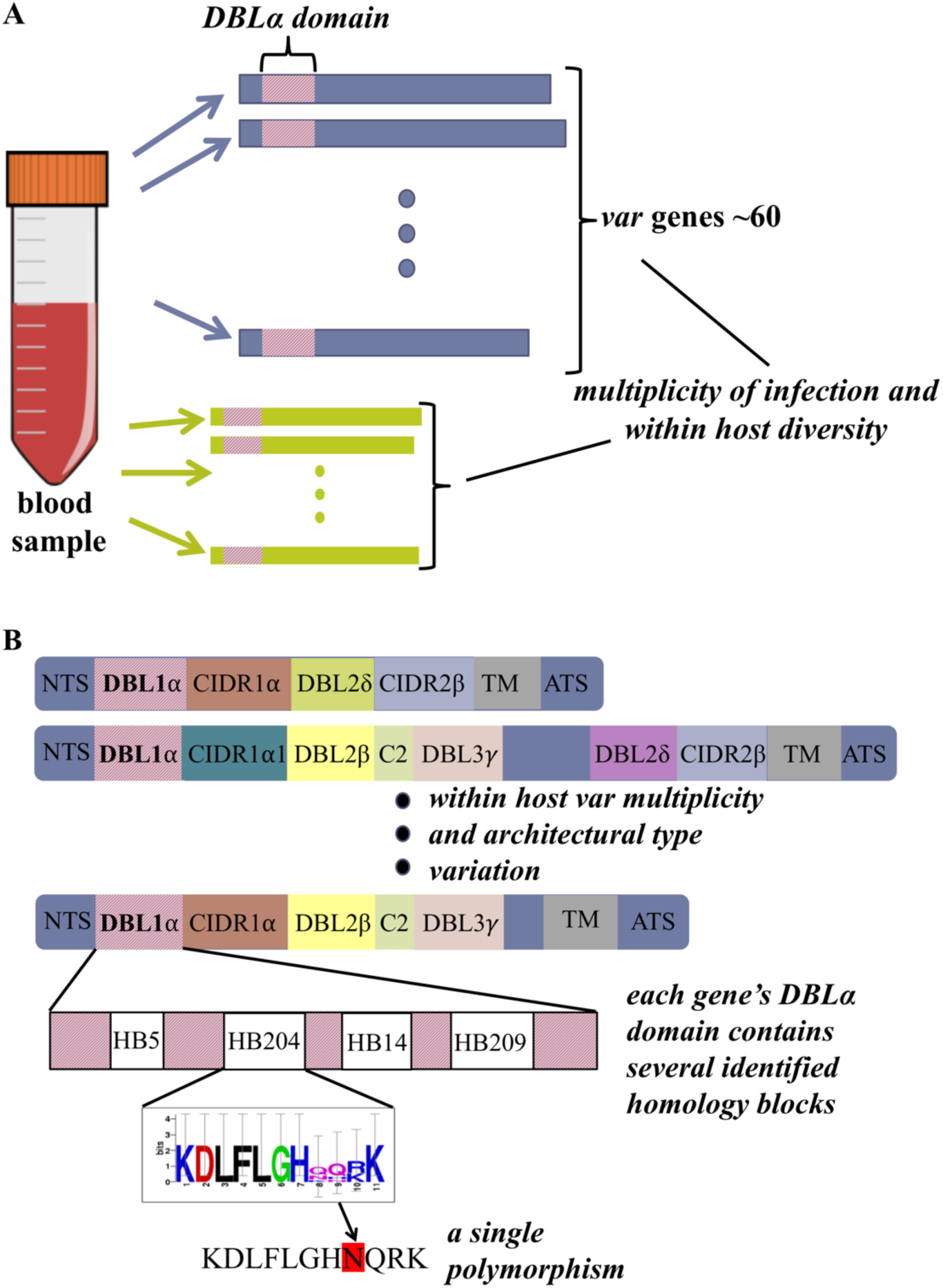
Amino-acid polymorphisms of PfEMP1. **A.** Each *Plasmodium falciparum* parasite contains within its genome a set of approximately 60 *var* genes. A set of *var* genes, representing one or more infections, is sequenced from each blood sample, taken from individuals participating in the pilot study. The *DBLa* domain, located in the head domain of most *var* genes is sequenced and further analyzed. **B.** PfEMP1 proteins can be classified into multiple architectural types with an alternative domain structure, three of which are depicted for illustration; N-terminal segment (NTS), DBL, CIDR, C2 domain, transmembrane (TM), acidic terminal segment (ATS) (38). The *DBLa* domain is further analyzed for the identification of short alignable sequences termed homology blocks. Sequences that compose homology blocks contain specific amino-acid motifs, but are not exact copies of each other. Variation, or single amino acid polymorphisms, within homology blocks is used in further analyses and is tested for association with disease clinical manifestation.

### Genomic samples

Three genomic isolates were used as positive controls for our sequencing and analysis methods: 3D7, DD2, and HB3. These isolates have a known multiplicity of infection (=1), and the number of *var* genes that exist per genome has been previously established. The sequence of each of the DBLα tag sequences within these genomes is also known with high accuracy. For these isolates we can distinguish sequencing errors from within-type variants, and therefore identify multiple *var* sequences of the same type.

### Transcription samples

The expressed sequences and the clinical data for 250 isolates were obtained from the online supplementary information of (21). The Warimwe et al. study included children recruited between August 2003 and September 2007 from the Kilifi District Hospital, situated at the coast of Kenya. In their study DBLα sequence tags were amplified from parasite cDNA sampled from each of 112 (44.8%) children with severe malaria, 105 (42%) children with symptomatic non-severe malaria, and 33 (13.2%) asymptomatic children.

Here, we divided the “expression dataset” of Warimwe et al. randomly into a validation dataset and a training dataset, containing 175 (70%) and 75 (30%) individuals, respectively. Training and normalization were performed using the training dataset expression levels only. Prediction is reported with respect to the validation dataset.

### Transformation of expression rates and rosetting level

Prior to performing all linear and logistic regression analyses, the expression rates of particular *var* types (i.e., cys2, A-like, group 1, group 2, group 3, BS1/CP6 and H3sub *var* genes), of homology blocks (i.e. for all 29 HBs), and of homology block polymorphisms (i.e. 2148 polymorphisms in the Warimwe et al. dataset) were normalized and log transformed. Normalization was performed by dividing with the corresponding median expression rates in non-severe and asymptomatic cases in the training dataset. Prior to the log transform, extreme (i.e. zero and one) expression values were replaced with 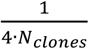 and 1 − 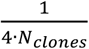, respectively. Rosetting and parasite density levels equal to zero were replaced with 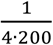 and 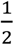, respectively, prior to the log transform. These types of replacements were also carried out by (45), and they are for numerical reasons and to account for the limited sensitivity of detection.,.

### Predictive Ability

Because of the large dimensionality of the data, and the large number of predictive variables in comparison to the number of study individuals (p>>n), the predictive ability of different models was evaluated using a separate dataset which was not included in data normalization or training. For binary-outcomes, predictive ability was calculated using the ‘area under the curve’ (AUC) statistic. Given a list of disease state prediction scores generated by the model for each individual in the validation dataset, the AUC is the probability that a disease positive individual will get a higher score in comparison to a disease-free individual. For continuous outcomes, predictive ability was calculated using the *forecast skill* (SS) in comparison to the training dataset mean. This is equivalent to an *r*^2^ statistic with the difference that the linear fit is created using the training dataset and the statistic is calculated for the validation dataset.

### Model Selection

Model selection was performed using the ‘Sparse Group LASSO’ R package (66). Similar to the lasso method, this method minimizes a weighted sum of the model prediction error and the absolute value of the regression coefficients. As such it prefers sparse solutions. A stochastic implementation of this procedure is repeated several times, the model used includes only variables which appear in at least a certain percent of models, referred to as stable variables. A threshold of 90% of models was used for logistic models. A different threshold of 20% evaluated empirically was used for inclusion of variables in the linear prediction of rosetting.

The sparse group lasso method assigns a penalty to groups of coefficients together. Polymorphisms were assigned to groups based on belonging to the same site in the homology block sequence, the remaining variables were not assigned into groups. An equal weight was given to the lasso and group lasso penalties (α=0.5).

### False discovery rates

False discovery rates (q-values) were calculated using the algorithm described in (67) R package 1.38.0. This method offers a more powerful way of correcting for multiple comparisons in relation to e.g. Bonferroni correction, and is expected to provide more robust estimates in comparison to Benjamini-Hochberg (68). It uses an interpolation approach for estimating the excess number of p-values of a given value and thus the false discovery rate.

## RESULTS

### Diversity of amino-acid polymorphisms in Ghana and in Kenya

We measured the fraction of individuals, and the fraction of *vars* within each individual, in which specific single amino-acid polymorphisms within homology blocks were present. This was done in genomic data from the study population in the Bongo district of Ghana in 2014, and in expression data from the Warimwe et al. 2009 study in the Kalifi district of Kenya.

We identified a total of 2701 single amino acid polymorphisms (PMs) occurring at 316 different sequence positions (sites) (Fig. 2). 2098 (77.6%) polymorphisms across 305 (97%) sites were shared between the two study populations/dataset types. A total of 48 (1.8%) PMs were only present in expression data from Kenya, whereas a higher total of 555 (20.5%) PMs were only present in the genomic dataset from Ghana. The fraction of individuals with a given PM (the population-level frequency) was highly correlated between the two studies (Spearman’s rank ρ=0.96, p=0) (Fig. 2), indicating that the population-level frequency of PMs is actively maintained by evolution across vast geographic ranges (the two study sites are 5000km apart), even in the diverse parasite populations of sub-Saharan Africa.

**Figure 2.**
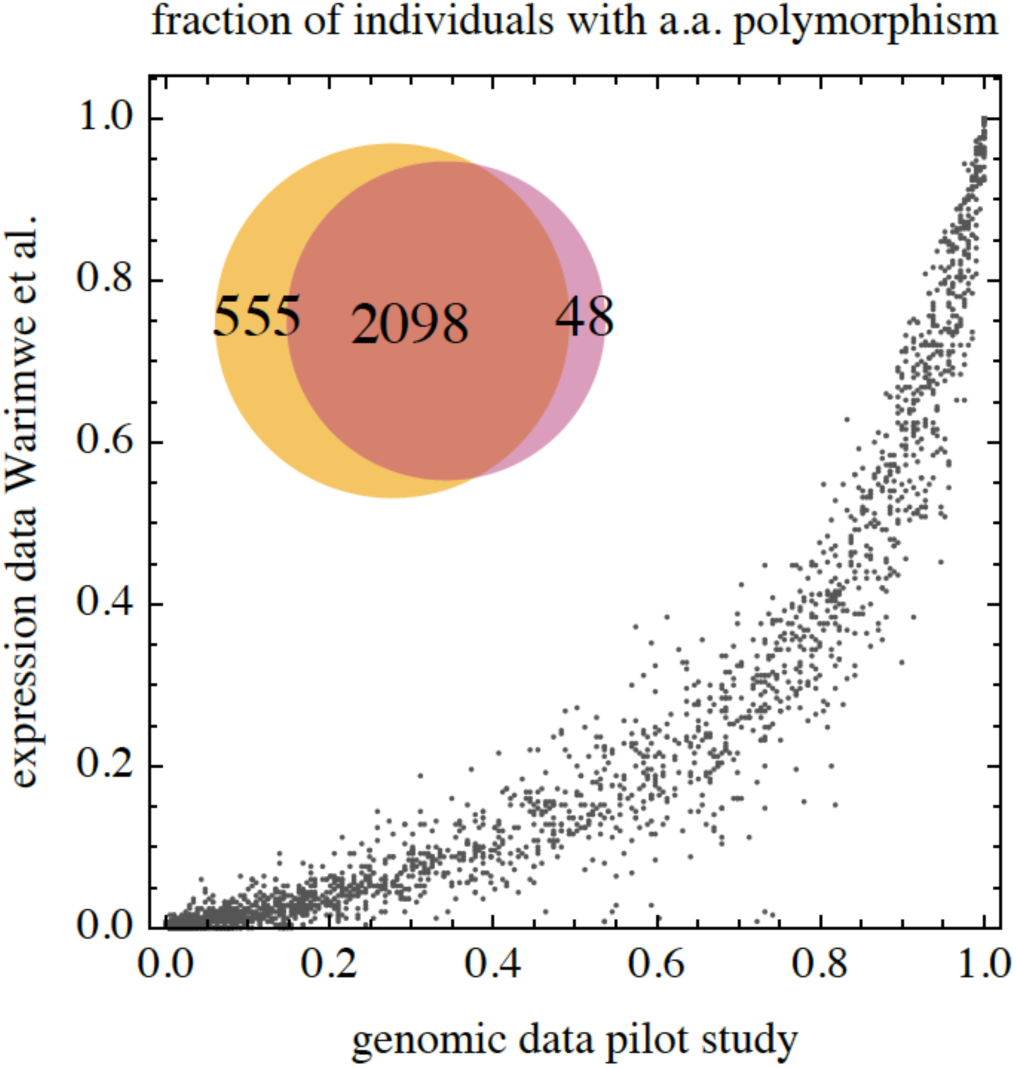
Frequency of amino-acid polymorphisms in the two study populations. The fraction of *vars* in which single amino-acid polymorphisms within homology blocks are present in the study population in Ghana, and in expression data from the Warimwe et al. study. A total of 2701 amino acid polymorphisms were identified in either genomic sequences in the Ghana study (left) or expression data in Kenya (right). 2098 amino-acid polymorphisms were shared between the two study populations, 555 were unique to genomic data in the Ghana study, and 48 were unique to the expression data in the Kenya study.

On average of 1030±290 (mean±std) PMs were identified per individual from the genomic data from Ghana, while roughly half (550±290 PMs) were identified per individual from the transcription data from Kenya. Because there are multiple *var* copies per parasite genome, and often multiple parasite genomes within a given individual host, PMs can occur multiple times within an individual. Therefore, PMs have an individual-level frequency in addition to a population-level frequency. The individual-level frequency of a given PM was correlated among the individuals within a population (average Spearman’s <ρ>=0.61±0.1 in Kenya, <ρ>=0.78±0.1 in Ghana) and between the two populations (<ρ>=0.60±0.1). These correlations indicate that, in addition to the population-level frequency, the individual-level frequency of PMs is conserved. Furthermore, in general, variation between individuals is greater with respect to expression data as compared to genomic data. The one main exception to a high degree of population-level conservation among PMs was that 11 unique polymorphic sites in HB190 were only identified in the genomic dataset from Ghana.

The conservation of the different PMs and their frequencies in the two study populations suggests that these PMs may be evolutionarily conserved over large geographic spaces; thus, associations observed between PMs and pathophysiology in geographically restricted populations, such as the two we examine here in Kenya and Ghana, may be relevant to larger populations, or possibly even globally. We therefore continue by identifying such associations and their relation to previous research findings in the literature. Unique to this study, we explore polymorphisms within HB5 and HB14, which are two common homology blocks present within most *var* genes. We identified 628 amino-acid PMs within them, 481 (76.6%) of which are present in both datasets.

### Association with severe disease and rosetting

We used the expression levels of the *classic var* types, of homology blocks, and of homology blocks polymorphisms to predict severe disease and rosetting. In addition, we evaluate the level of parsitemia both as an independent variable and as an interaction term. *Classic var* types were defined by the presence/absence of specific motifs in the case of cys2PoLV groups and h3sub *var* types, and by network analysis in the case of A-like and BS1/CP6 *var* types, and were obtained from the online supplementary information of (21). The expression levels of *classic var* types and of homology blocks were used in previous studies to predict severe disease (13, 21). A model selection scheme was employed to identify the best logistic model using two initial datasets considering either the expression of classic *var* gene groups or the expression of homology blocks and their polymorphisms. In both cases, we found that an interaction term with parasitemia was included in the optimal model.

Predictive ability was measured using an independent subset of the dataset which was randomly selected and was not used in training or data normalization. Prediction of severe disease based on gene expression data was good (AUC=0.82 for *vars*, 0.83 for homology blocks and their polymorphisms) and was not significantly different when considering the two alternative datasets (Fig. 3). However, the use of classic *var* subtypes included fewer variables namely grpA *var* genes, the interaction term of the H3 *var* subset with parasitemia and parasitemia levels (Table 1). The inclusion of age as a predictor did not improve on the prediction of severe disease.

**Figure 3.**
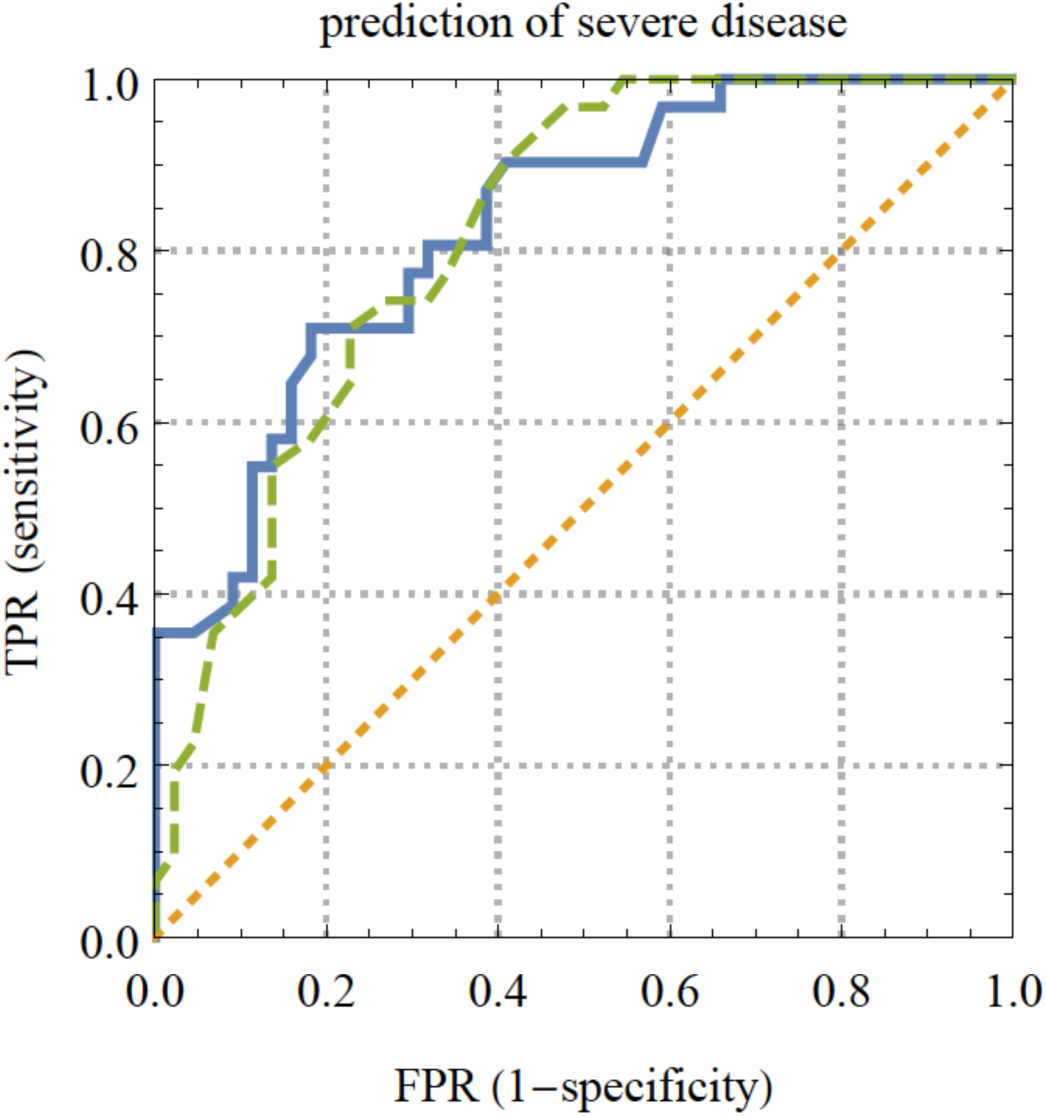
ROC curve for prediction of severe disease in an out of bag dataset. The area under the curve represents the probability that a randomly chosen subject with severe disease is correctly rates or ranked with greater suspicion than a randomly chosen individual without severe disease. sold line = prediction based on amino acid polymorphisms, dashed line = prediction based on *“classic” var* gene groups. Scores were calculated using a logistic regression model, and ROC curves represent prediction in an out of bag dataset of 75 individuals not used in the normalization of expression levels or in the training of the model.

**Table 1.** Statistics for logistic regression models predicting severe disease

| model data | stable predictive variables | AUC |
| --- | --- | --- |
| **HBs, PMs,**<br>**interaction terms**<br>**with parasite**<br>**density (pdt)** | **HB60, HB204, HB219, HB345, HB367,**<br>HB163×pdt, HB171×pdt, HB219×pdt,<br>HB14<sub>19I, K, M</sub>,<br>**HB60<sub>6D</sub>×pdt, HB60<sub>7S</sub>×pdt, HB60<sub>10Q, R</sub>×pdt,**<br>HB64<sub>6Q, R, V</sub>×pdt,<br>HB204<sub>8S</sub>×pdt, **HB204<sub>9E</sub>×pdt, HB204<sub>10E</sub>×pdt, HB402<sub>1K</sub>×pdt,**<br>**pdt** | **0.83** |
| **classic *var* genes and**<br>**interaction with**<br>**parasite density** | **grpA, H3×pdt, pdt** | **0.82** |

positive effect independent variables are shown in **boldface**
subscripts indicate the position and a polymorphism and the amino-acid variant

When predicting rosetting, *forecast skill* (SS) was considerable for *vars* (SS=0.41) and when prediction was made based on homology blocks and their polymorphisms (SS=0.43) (Fig. 4). As was the case for predicting severe disease, there was no significant difference between the predictive ability of the two models, however the model which included the classic *var* types had fewer variables. Interestingly, the prediction of *rosetting* and severe disease included the same two *var* subsets: *grpA* and *H3*, however the rosetting model included *grpA* rather than the *H3* interaction term with parasitemia (Table 2).

**Figure 4.**
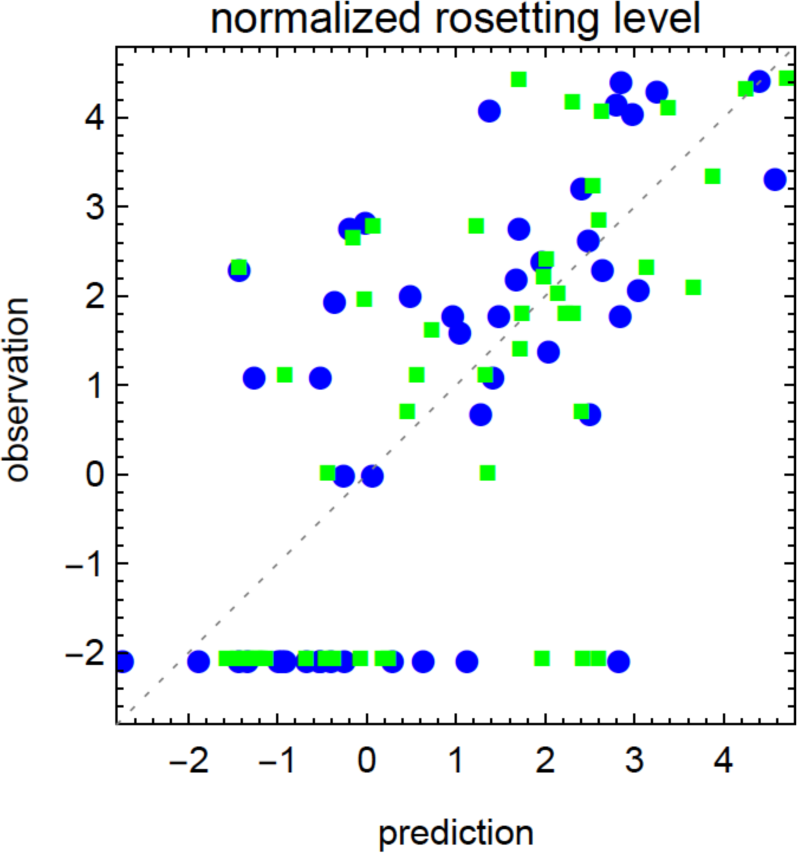
Prediction of rosetting. Prediction of normalized rosetting levels using a linear model and the transcription levels of *var* genes or of HBs and their amino acid polymorphisms. Prediction is shown for an out of bag dataset of 75 individuals not used in the normalization of expression levels or in the training of the model. • = prediction based on amino acid polymorphisms, ▪ = prediction based on *“classic” var* gene groups.

**Table 2.** Statistics for linear regression models predicting rosetting

| model data | stable predictive variables | SS |
| --- | --- | --- |
| **HBs, PMs,**<br>**interaction terms**<br>**with parasite**<br>**density (pdt)** | HB5<sub>7 L,M, I</sub>,<br>HB14<sub>22 Q, R, T, Y</sub>, HB14<sub>23 C, D, E, H</sub>,<br>HB54<sub>14 K, L</sub>,<br>HB60<sub>11 G, N, P, Q, S, V</sub>, HB60<sub>12 H, K, Y</sub>, HB60<sub>13 C</sub>, HB60<sub>14 O</sub>,<br>HB210<sub>1 K</sub>, HB210<sub>2 A, E, G, Q, T</sub>, HB210<sub>3 R</sub>, HB210<sub>4 Y</sub>,<br>HB210<sub>5 E, G, Q, R</sub>, HB210<sub>9 Y, F</sub>, HB210<sub>10 L, Y</sub>, HB210<sub>11 F, L, Y</sub>,<br>HB210<sub>12 K, Q</sub>, HB210<sub>13 L, Y</sub>,<br>HB219<sub>1 K, R</sub>, HB219<sub>4 F</sub>, HB219<sub>6 P, R</sub>,<br>HB14<sub>21 R, S</sub>×pdt,<br>HB36<sub>30 G, N, K, L</sub>×pdt,<br>HB60<sub>10 D, E, G, H, I, S, T, W</sub>×pdt, HB60<sub>11 O, C, G</sub>×pdt,<br>HB60<sub>20 O, A, D, P</sub>×pdt, HB60<sub>21 H, I, L, P, Q, S</sub>×pdt, HB60<sub>22 P, T</sub>×pdt,<br>HB60<sub>23 N, T, V</sub>×pdt, HB60<sub>24 I, L</sub>×pdt,<br>HB64<sub>2 T</sub>×pdt, HB64<sub>8 A</sub>×pdt, HB64<sub>9 K, N, R, S, Y</sub>×pdt,<br>HB79<sub>12 A, D, E, G, K</sub>×pdt,<br>HB131<sub>5 Q, R</sub>×pdt, HB131<sub>6 A</sub>×pdt,<br>HB204<sub>8 S</sub>×pdt, HB204<sub>9 E</sub>×pdt, HB204<sub>10 E</sub>×pdt,<br>HB260<sub>20 Q</sub>×pdt, HB260<sub>23 N</sub>×pdt,<br>HB345<sub>8 Y</sub>×pdt, HB345<sub>9 F, L, Y</sub>×pdt,<br>HB402<sub>7 T</sub>×pdt,<br>HB210<sub>13 L, Y</sub>×pdt,<br>HB367<sub>5 N</sub>×pdt, HB367<sub>6 F, Y</sub>×pdt, HB367<sub>7 F</sub>×pdt, HB367<sub>8 R</sub>×pdt,<br>HB367<sub>9 K, P</sub>×pdt<br>pdt | **0.4** |
| **classic *var* genes and**<br>**interaction with**<br>**parasite density**<br>**(pdt)** | grpA, H3, grpA×pdt, pdt | **0.4** |

positive effect independent variables are shown in **boldface**
subscripts indicate the position and a polymorphism and the amino-acid variant

We continued by testing whether specific polymorphisms are associated with the *grpA* or *H3 var* subsets (Table 3). Using a similar model selection scheme, we find a set of specific polymorphisms and homology blocks with expression levels correlated with the expression of *grpA vars* (*forecast skill* SS=0.93, p=0) (Fig. 5A). To a lesser degree, a different group of polymorphisms is associated with *H3 vars* (SS=0.20, p=10^−8^) (Fig. 5B). Based on these results it is likely that a substantial portion of the ability of PMs to predict severe disease relies on their association with grpA-like *vars*.

**Table 3.** Statistics for logistic regression models association with *grpA* and *H3 var* subsets

| predicting transcription of | stable predictive variables | SS |
| --- | --- | --- |
| *grpA* | **HB60**, **HB179**, **HB219**, **HB367**, **HB402**, **HB486**,<br>HB5<sub>20</sub>S, **HB14**<sub>1</sub>V, **HB14**<sub>4</sub> A, G, N | **0.93** |
| *H3* | HB219, **HB402**, **HB219**<sub>8</sub> D, K, Q, **HB219**<sub>9</sub> A,D,E,K,Q,<br>**HB219**<sub>11</sub> A, K, **HB5**<sub>7</sub> L, M | **0.20** |

positive effect independent variables are shown in **boldface**
subscripts indicate the position and a polymorphism and the amino-acid variant

**Figure 5.**
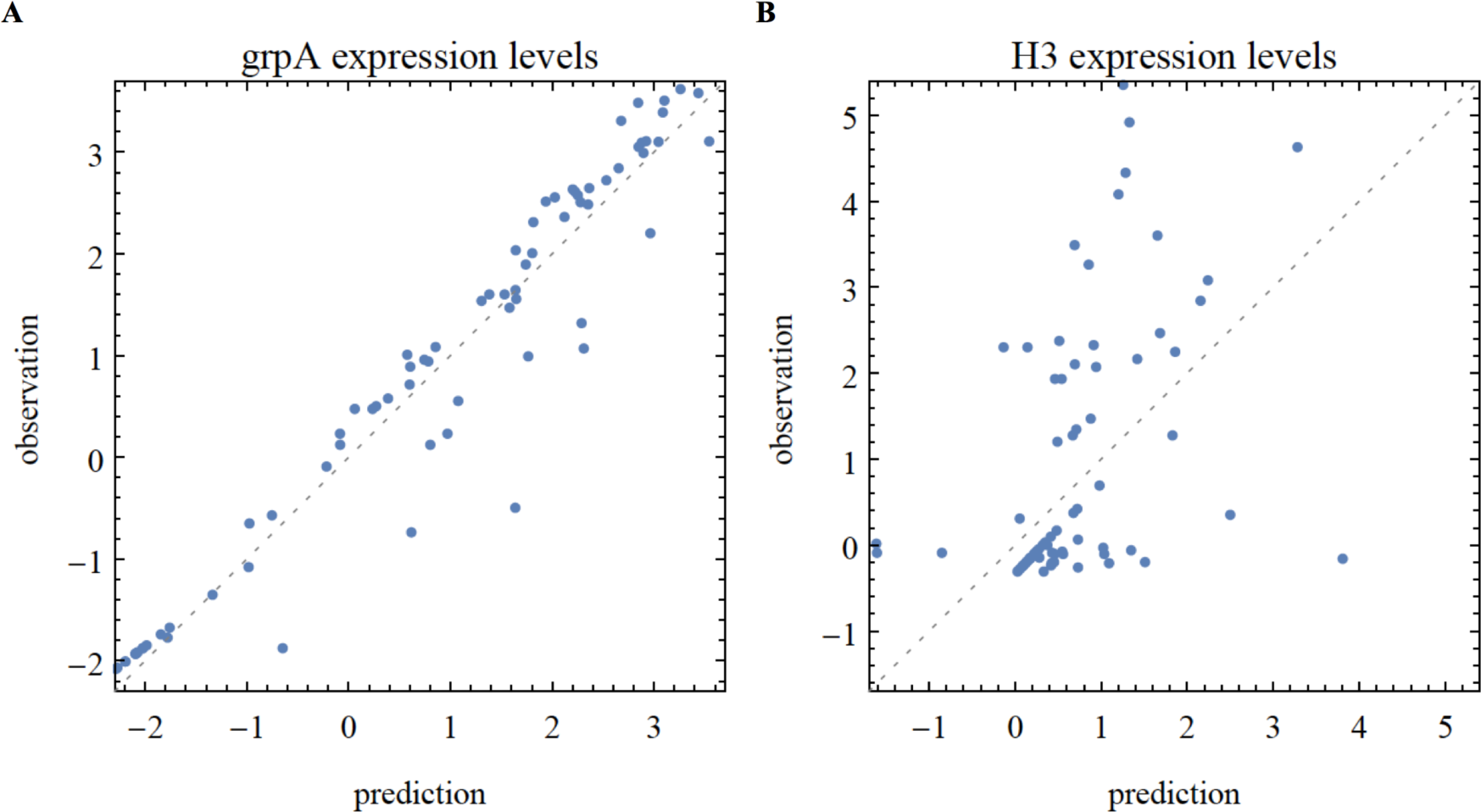
Association between group A like and H3 *vars*, homology blocks and their polymorphisms. Prediction is shown for an out of bag dataset of 75 individuals not used in the normalization of expression levels or in the training of the model. **A** Prediction of the transcription levels of *group A like vars* using a linear model and the transcription levels of HBs and their amino acid polymorphisms. **B** Prediction of the transcription levels of *H3 vars* using a linear model and the transcription levels of HBs and their amino acid polymorphisms.

### Association of *var* gene polymorphism with age and parasitemia

In the Warimwe et al. study (45), the overall number of *var* gene clones declines with age (ρ=-0.26, p=0.00019). We tested whether specific homology blocks, or homology block polymorphisms, had changes in their transcription levels in relation to the total number of clones, in a way which go beyond the overall decline with age. We calculated the p-value of the linear regression models (Fig. 6) and corrected for multiple testing as described in (69). We found HB60 to have a moderately lower relative expression rate with increasing age (ρ=-0.30, p=1.2·10^−8^, FDR=4.6·10^−5^) while HB36 displayed an increased relative expression rate with increasing age, however this relationship was not significant (ρ=0.16, p=0.0017, FDR=0.18). HB36 still maintained an overall decline in transcription with age. Similarly, we calculated the association of HB polymorphisms with age. We found that the expression rates of four of HB60s polymorphisms had a similarly strong and declining expression level with host age however none showed a stronger correlation than the expression of the HB itself.

**Figure 6.**
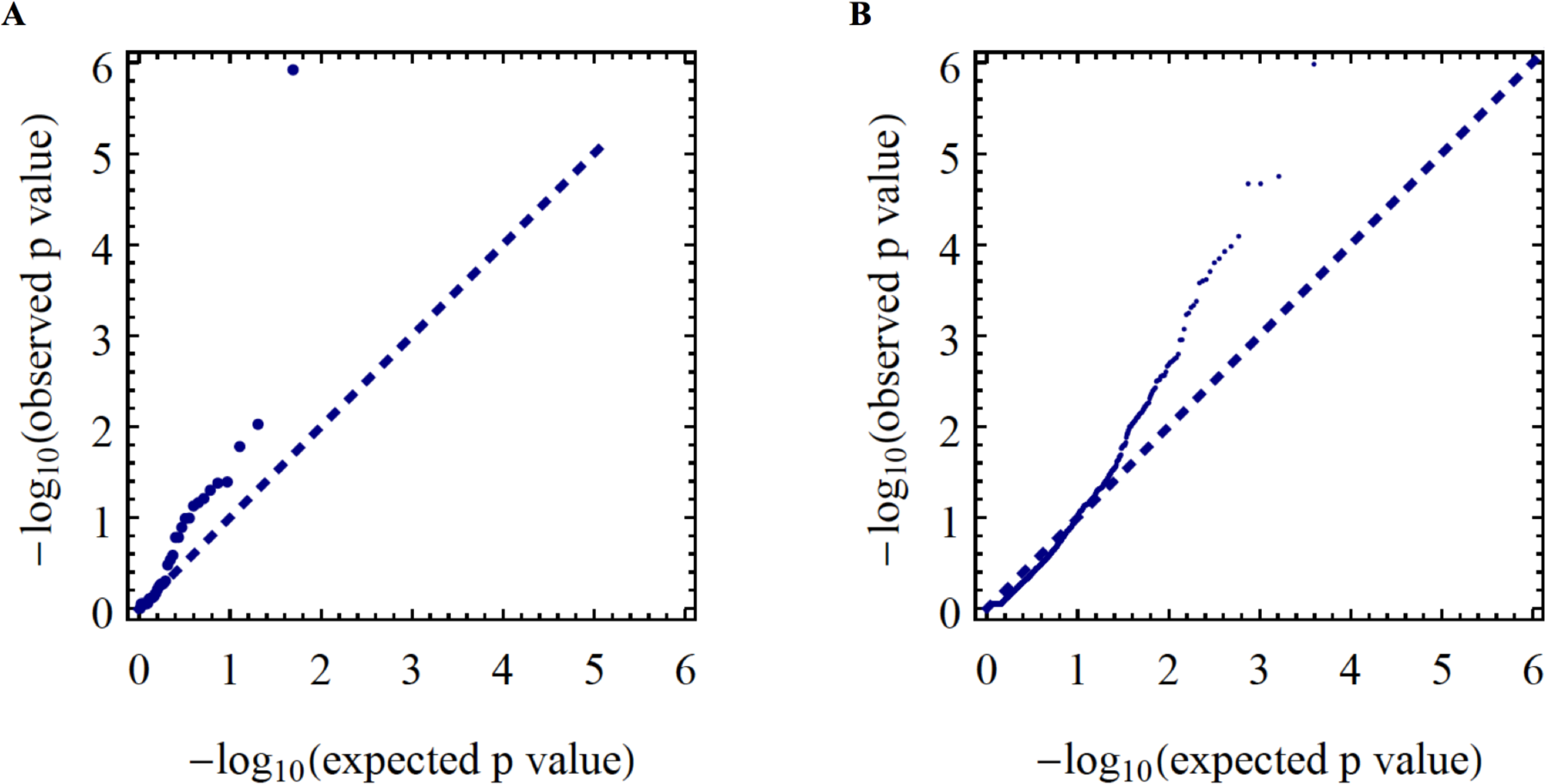
Q-Q plot for association between HB and HB polymorphisms with age. The quantile-quantile (Q-Q) plot showing the significance of linear models associating age with the relative expression of HBs (**A**) or of HB polymorphisms (**B**).

Finally, we tested whether *var* types and/or *var* HB polymorphism were associated with parasitemia. Neither the relative expression of *var* genes nor the relative expression of homology blocks was associated with parasitemia directly. An association between polymorphisms and parasitemia was only detectable in an ensemble of selected logistic models (*Pearson’s* ρ=0.36, p=0.02) and not in a consensus model including only stable selected variables.

## DISCUSSION

Antigenic diversity in *P. falciparum* is a major obstacle for developing vaccines against malaria. The identification of functional and conserved antigenic targets involved in pathogen virulence is greatly needed. The high diversity of the *var* family has made the identification of reliable *var* single nucleotide polymorphisms (SNPs) difficult (70). In our study, we limit our analysis to amino-acid level variation within homology blocks in the *DBLα* domain with the aim of identifying reliable variation of functional consequence. We identify conserved amino-acid polymorphisms within the DBLα domain of *var* genes that are associated with severe pathophysiology and with rosetting. Polymorphisms are a part of multiple conserved and functional protein domain sites and prediction of severe disease based on their transcription level compares in its skill to the prediction of severe disease based on the expression levels of classic *var* groups. However, amino-acid polymorphisms within homology blocks can uniquely illuminate additional and potentially more direct relationships between protein variation, immunity and function.

We observe the great majority of homology block polymorphisms in both the Kenyan and Ghanaian populations, indicating their evolutionary conservation across large distances of time and space. Furthermore, the fraction of *vars* with a specific polymorphism was correlated between pairs of individuals sampled from the two populations (mean Spearman’s <ρ>=0.60) and within each population (<ρ>=0.61±0.1 in Kenya, <ρ>=0.78±0.1 in Ghana). The two study populations are located in very different geographic locations (East versus West Africa), they were sampled more than five years apart, and they represent distinct collection methods (expression versus genomic data). Together, our findings suggest that there is a conserved set of *var* polymorphisms at the genomic level that is manifest in proportional levels of transcription. It also suggests that variation within homology blocks is conserved and maintained.

The maintenance of homology block polymorphisms across populations in light of the high rates of ectopic recombination known to characterize the *var* family (71–73), suggests that balancing selection may be at play. Balancing selection may also be responsible for maintaining similar frequencies among variants across populations if certain ratios between alternative PMs at a given site are adaptively optimal for the parasite. Alternatively, conserved frequencies among variants may be attributable to the existence of recombination constraints (or so-called *var* recombination hierarchies) that orchestrate recombination preferentially among certain *var* types and/or maintain ratios of certain *var* types within genomes. They may also be due to shared, recent evolutionary history—i.e. the neutral inheritance and conservation of similar frequencies of variants. The main *var* groupings reflect both sequence and functional divergence, so it is possible that homology block polymorphisms reflect this functional divergence as well.

Consistent with findings in literature, and with findings based on the same primary data, we found that severe disease correlated with the expression of specific ‘classic’ *var* gene types and with higher parasitemia (21, 47, 74). The optimal model for predicting severe disease that we present here differs from *Warimwe et al.* in that it includes only *group A var*s, the interaction term of the *H3* subset with parasitemia, and parasitemia levels as an independent variable (Table 1, Fig. 3). In contrast with other studies, we find no association between group B *vars* and severe disease was identified (17, 44). The dependence on the expression levels of *vars* and on the interaction term with parasitemia may help explain why control of parasitemia by itself is insufficient in preventing severe disease (9).

Interestingly the same *var* subgroups were also correlated with the rosetting phenotype (Table 2), although this relationship included a different interaction term with parasitemia. This is consistent with the notion of rosetting as a factor contributing to virulence (57). The prediction of rosetting based on the expression of these classic *var* groups and their interaction with parasitemia explained 40% of the variance in the validation dataset (Fig. 4). Additional information regarding the host genotype, specifically relating to ABO blood groups may improve prediction of severe disease and of rosetting (36, 57).

Also consistent with findings in literature, we find an association between certain homology blocks and severe disease, specifically, the transcription of HB60, HB204, HB219, HB367 was associated with severe disease (13). The transcription of HB345, previously associated with a milder symptom gene group, was instead associated with severe disease in the models we describe here. Consistent with previous work, we found that the transcription of HB163 and HB171 was associated with milder symptoms, however in a form including an interaction term with parasitemia. The transcription of HB219, positively associated with severe disease, was negatively associated with severe disease in its interaction with parasitemia.

Adding to the inquiries of previous studies, we also consider variation within homology blocks as predictive variations for disease symptoms. For instance, a polymorphism of HB402: HB402_1K_, was associated with severe disease in its interaction with parasitemia (Table 1, Fig. 3). The HB402 motif itself was previously associated with severe disease, while not considering individual variation (13). HB5 and HB14 are two common homology blocks present within most *var* genes, and we identified 628 amino-acid PMs within these two HBs, the majority shared between the two datasets. Thus, our findings suggest that variation within common homology blocks, such as HB5 and HB14, should also be considered when assigning functional, virulence, immunogenic or evolutionary attributes to *var* types. For example, the transcripts of three variants of HB14 at amino acid position 19 (I, K, M) had either positive or negative associations with severe disease (Table 1).

Since the predictive ability of HBs and HB polymorphisms did not surpass prediction based on ‘classic’ *var* genes, we measured the association between the different polymorphisms and the predictive grpA-like and H3 subset *vars*. We identify a strong association between the expression of six previously severe implicated HBs (HB60, HB179, HB219, HB367, HB402, HB486) and grpA-like *vars* (Table 3). In addition, we find novel associations between polymorphisms of HB14 and HB5 and grpA-like *vars* (Table 3). A weaker association between polymorphisms and the H3 subset was identified and includes: HB219 and its polymorphisms, the polymorphisms of HB5, and HB402 (Table 3). Several HB5 polymorphisms were associated with group A-like and H3 *vars* but not with severe disease. Such variation is of interest since it may shed light on the distinction between high and low virulence grpA-like *vars* (Table 1, Table 3).

The expression of group A-like *vars* is known to be associated with young age and a low degree of acquired immunity (75), which may provide protection against severe disease. An alternative explanation to the pattern is that group A *vars* may be expressed later during an infection. An expression hierarchy such as this, coupled with higher parasite clearance rates in older individuals, may also explain the phenomenon. However, it does not fit with the presence of chronic asymptomatic infections in adults (76). In our study, HB60 is associated with grpA-like *vars* and with severe disease, while in previous studies HB36 was shown to be associated with milder symptoms (13). In our analysis, we found that HB60 had moderately but significantly lower relative expression rates in older individuals (ρ=-0.30, p=1.2·10^−8^, FDR=4.6·10^−5^), while HB36 had higher relative expression rates in older individuals (ρ=0.16, p =0.0017, FDR=0.18) (Fig. 6). Both hypotheses for the correlation with age—an expression hierarchy or specific immunity—could generate the observed patterns, so we view both as plausible in this case.

In the prediction of both severe disease and rosetting based on homology block polymorphisms, the majority of associations we identify are not significant individually from the standpoint of classic logistic or linear regression, but rather provide good predictive ability as an ensemble. Larger sample sizes or a meta-analysis approach may help attain statistical significance at the finer-grain level of detail of individual HB polymorphisms. Alternatively, an experimental approach focusing on specific polymorphisms or homology blocks such as HB60 or HB219 would provide more data of this type. Additional limitations to this work include the fact that the dataset sampled from Ghana did not include any *var* expression level data, and that there was no sampling of asymptomatic individuals in the Kenyan dataset. In addition, the conserved frequency distribution of polymorphisms shared among the study sites may reflect *P. falciparum* populations in Africa, rather than globally. Limited sample size significantly impacted our ability to verify specific associations between polymorphic site variation and pathophysiology.

The association of specific homology blocks and polymorphic sites within homology blocks with severe disease further emphasizes the potential for developing vaccines that could target genes responsible for severe malaria, and thus reduce the burden of mortality and morbidity associated with *P. falciparum*. Ideally such vaccines will protect young children in Africa against severe disease in wide geographical areas and will not lose their effectiveness through parasite evolution (e.g., antigenic drift). Consistent with previous findings, such putative vaccines could include parasite *var* gene epitopes involved in severe pathophysiology. Our findings suggest that multiple yet conserved sequence motifs are associated with severe disease, and as such the acquisition of multiple antibodies against them may be sufficient to gain protection. In the case of placental malaria, protective antibodies in pregnant women have been shown to cross-react across geographically diverse placental isolates (77, 78). This cross-reactivity has served to motivate the ongoing work on vaccines for pregnant women (77). Targeting variation involving conserved sites within grpA var genes could be of equally great benefit.

## ACKNOWLEDGMENTS

We wish to thank the participants, communities and the Ghana Health Service in Bongo District Ghana for their willingness to participate in this study. We would like to thank the field and teams for their technical assistance in the field, as wells the laboratory and research personnel at the Navrongo Health Research Centre for sample collection and parasitological assessment/expertise. Additionally, we would like to thank laboratory personnel at New York University, Noguchi Memorial Institute for Medical Research, and The University of Melbourne for their assistance with laboratory experiments. Finally, we would like to acknowledge Michael Duffy, for his helpful input related to this work. MP is an Investigator at the Howard Hughes Medical Institute.

## FINANCIAL SUPPORT

This research was supported by the Fogarty International Center at National Institutes of Health [Program on the Ecology and Evolution of Infectious Diseases (EEID), Grant number: R01-TW009670]; and the National Institute of Allergy and Infectious Disease, National Institutes of Health [Grant number: R01-AI084156].

